# MORE interpretable multi-omic regulatory networks to characterize phenotypes

**DOI:** 10.1101/2024.01.25.577162

**Authors:** Maider Aguerralde-Martin, Mónica Clemente-Císcar, Ana Conesa, Sonia Tarazona

## Abstract

The identification of phenotype-specific regulatory mechanisms is crucial for understanding the molecular basis of diseases and other complex traits. However, the lack of tools capable of constructing multi-omic, condition-specific regulatory networks remains a significant limitation. He re, we introduce MO RE (Multi-Omics Regulation), a novel R package for the inference and comparison of multi-modal regulatory networks publicly available at https://github.com/BiostatOmics/MORE. MORE supports any number and type of omics layers, integrates prior regulatory knowledge, and employs advanced regression-based modelling and variable selection techniques to identify significant regulators of target features.

We evaluated MORE on simulated datasets and benchmarked it against state-of-the-art tools. Our tool exhibited superior accuracy in identifying key regulators, model goodness-of-fit, and computational efficiency. Additionally, we applied MORE to an ovarian cancer dataset to uncover tumour subtype-specific regulatory mechanisms associated with distinct survival outcomes.

By providing a comprehensive and user-friendly framework for constructing phenotype-specific regulatory networks, MORE addresses a critical gap in the field of multi-omics data integration. Its versatility and effectiveness make it a valuable resource for advancing our understanding of complex molecular interactions and regulatory systems.

## Introduction

Unveiling the regulatory mechanisms driving disease or other phenotypes of interest in a biological system is currently one of systems biology’s most important goals and challenges. Many advances have been made in this field, from gene co-expression networks derived from transcriptomic data to multi-layer regulatory networks leveraging different omic perspectives [1]. Interestingly, single-cell technology has recently emerged as an unprecedented opportunity to study cell type-specific regulation. However, it is yet to be possible to generate regulatory models encompassing a wide variety of omic layers since no more than two omic modalities can be measured on the same cell, and, in addition, few omic combinations are commercially available [2]. Therefore, bulk multi-omics is still the only option to model complex regulatory mechanisms comprehensibly.

Although gene co-expression networks can provide valuable information about the biological system under study and many methods have been developed to this end [3, 4, 5, 6, 7, 8], this information is limited by the fact that gene expression data are only considered. Incorporating other regulatory layers into the model is crucial to better understanding the complex molecular mechanisms triggered by disease or other environmental factors. Therefore, a multi-omics approach is pivotal to unravelling intricate regulatory mechanisms and providing a more holistic and interdependent view of biological processes. The increasing availability of multi-omics experiments makes it feasible, but efficient and versatile tools are required to infer such multi-layer regulatory models.

Many efforts have been devoted to developing efficient multi-omic regulatory networks (MO-RN) inference methods. However, there is still room for improvement. Versatile MO-RN methodologies must consider the heterogeneity of the data across omic technologies, with specific noise related to each modality, and the high dimensionality of the problem when combining omic features from several omic data types. Strategies to compare MO-RN across two or more phenotypes are also required. Finally, the biological interpretation of the multi-modal regulatory models obtained is still a bottleneck in this type of analysis [9, 10] that must be addressed. Moreover, some additional difficulties may be faced: (i) some methods can only analyse a limited number of omics (two or three at most); (ii) other methods are designed to study regulation between specific omic modalities (e.g., gene expression and methylation, gene expression and miRNA expression, etc.); (iii) some methods can computationally handle a reduced number of regulations and features; and (iv) some tools are no longer available because they could not be maintained by the research groups [11]. Next, we review some examples of these constrained yet valuable tools to highlight the existing gaps and underscore the critical need for novel integrative methodologies to infer MO-RN.

While the number of modalities measured in multi-omic experiments keeps increasing to uncover a wider diversity of molecular aspects of the biological system, many existing MO-RN tools are limited to the type or number of omic modalities, making them unsuitable for analysing such datasets. Some examples are DCGRN [12] and BMNPGRN [13], both designed for modelling the regulation of gene expression from DNA methylation and Copy Number Variation (CNV) data by employing a LASSO approach on multiple linear regression models to identify the most relevant regulators. Moreover, these two methods are not publicly available anymore. Other tools, even if extensible to more than two omics, are limited to their use on specific modalities. MoNET [14] is an R GitHub package that creates MO-RN with single nucleotide polymorphisms (SNPs), genes or proteins and metabolites from a protein-protein interaction (PPI) network. TFmiR [15] was a web server created to study regulatory interactions between transcription factors (TFs), microRNAs (miRNAs) and target genes involved in disease pathogenesis. The SAMNetWeb webserver [16] combines gene expression and proteomic data to generate condition-specific networks based on functional enrichment analysis. IntOmics [17], a deprecated Bioconductor R package, was designed to integrate gene expression, CNV, DNA methylation and biological prior knowledge on Bayesian networks. The RACER [18] pipeline is based on regression models to predict gene expression from CNV, DNA methylation, transcription factors and microRNAs, and it is not provided as a standalone package. The iDINGO R/CRAN package [19] generates MO-RN based on chained directed and undirected graph models and renders differential networks. However, due to potential computational limitations, the developers do not recommend using it with more than a few hundred features, which requires a very astringent preliminary feature filtering on omic datasets. In addition, up to three omics are admitted, although any omic modality is accepted.

Although not explicitly designed for generating MO-RN, multi-omic methods based on dimension reduction strategies have become very popular in finding common patterns between omic modalities and features of each modality. MOFA [20] and mixOmics [21] are two examples of such tools. In contrast to the methodologies described above, these tools usually admit any number or type of omic modalities and are helpful for visualising relationships among omics, features, or samples. However, they lack the functionality of providing specific feature-based MO-RN for each phenotype or condition under study, and neither can incorporate prior biological knowledge into the models. For instance, the MINT option in the mixOmics R/Bioconductor package, one of the most popular multi-omics approaches currently used for multi-omics data integration, applies either single or multi-block Partial Least Squares (PLS) regression models with a Lasso-based approach for variable selection and provides correlation networks for the selected omic features. Nevertheless, these networks are not condition-specific. Moreover, prior regulatory information cannot be given to the model, which may result in discovering novel regulations that could be biologically unmeaningful.

Interestingly, we found two methods that provide condition-specific MO-RNs and are flexible regarding the number or types of omics. CANTARE [22] is an R pipeline that fits pairwise regression models between each pair of omics to build the network and then applies logistic regression on the highly connected subnetworks to compare conditions. CANTARE does not admit prior knowledge about potential regulations. Therefore, although de novo regulations could be found with this method, false regulatory relationships are likely to be inferred. The other tool is KiMONo [23, 24], which builds MO-RNs from prior regulatory knowledge. KiMONo applies a sparse group LASSO penalisation on a multivariate regression model fitted separately for each gene to obtain the key regulators of gene expression from multi-omics measurements. The fitted models are aggregated in the final heterogeneous multi-omics network.

Given the limitations or unavailability of most of the published MO-RN tools, we introduce the MORE *R* package for MO-RN inference. MORE (Multi-Omics REgulation) is, to the best of our knowledge, one of the first software tools capable of generating condition-specific regulatory networks for any number or types of omic data, admits prior biological knowledge about regulations, and provides functionalities for comparing the resulting networks and guiding the biological interpretation of regulatory relationships. MORE relies on a regression framework and advanced variable selection techniques to identify global and condition-specific significant regulators. Figure 1 provides an overview of the MORE methodology.

**Fig 1.**
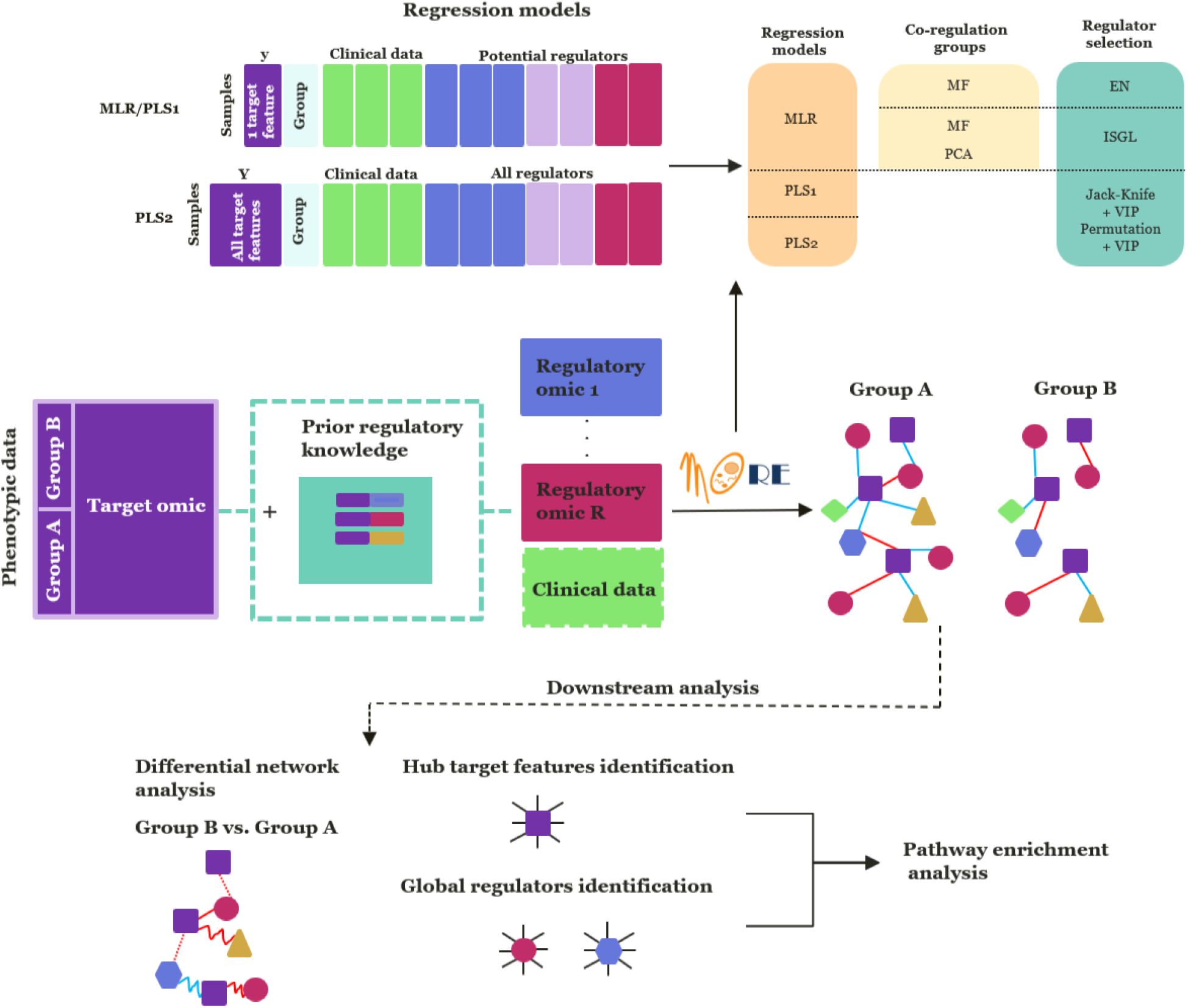
Overview of the MORE R package. MORE takes as input a target or “regulated” omic (e.g. gene expression), the regulatory omics (e.g. miRNA expression, DNA methylation, etc.) or clinical data, and the phenotypic groups to be compared (e.g. healthy/disease). Optionally, prior knowledge about regulatory relationships can also be provided (e.g. miRNA target genes). MORE applies regression models and advanced variable selection techniques to obtain the phenotype-specific regulatory networks. MORE networks display activation (blue) and repression (red) regulatory relationships, and the functionalities for network biological interpretation include differential network analysis to compare phenotypes, identification of hub target features and global regulators, and functional enrichment analysis of sets of relevant features in the regulatory network.

## Methodology

### MORE general framework

MORE infers a different MO-RN for each biological condition or phenotype under study (e.g. control/disease, treatments, or stages of a particular disease) using a regression framework. For fitting the regression models, MORE requires a target omic with the “regulated” features (e.g., gene expression) and one or more regulatory omics, that is, the potential regulators (e.g., methylation, miRNA expression, chromatin accessibility, etc.), all of them measured on the same n patients or samples. Any omic modality is admitted, but let gene expression be the target omic for simplicity. Optionally, prior information about the potential target genes of each omic regulator can be provided. If not, MORE will consider all regulators as potential regulators of each gene.

The MORE strategy consists of fitting a regression model per gene, where the potential regulators of that gene will be the predictors in the model. As the regulatory mechanisms might differ for each phenotype or condition, such conditions and their interactions with the potential regulators will be included in the model. Again, for simplicity, let us consider just two conditions (e.g. control and disease), although the model can be applied to any number of conditions.

Let **y**_*g*_ be the expression vector of gene g (g = 1, …, G) in the n samples; R_*g*_ the number of potential regulators of gene 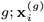 the values of the potential regulator i of gene g for the n samples, and **z** the dummy variable indicating whether a sample belongs to the control or disease group. Equation 1 describes the regression model to be fitted by MORE:

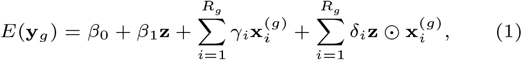

where ⊙ represents the Hadamard product, and β_0_, β_1_, γ_*i*_, and δ_*i*_ are the regression model coefficients. Note that when prior knowledge about potential regulations is provided to MORE, each gene may have a different set of regulators. Consequently, this initial model equation for each gene will be different.

MORE relies on Multiple Linear Regression (MLR) or Partial Least Squares regression (PLS) [25] models to fit the regression model presented in Equation 1. In the PLS case, there are two options: fitting a PLS1 model per gene where the response matrix **Y** is the vector **y**_*g*_ or a unique PLS2 model where **Y** = [**y**_1_, …, **y**_*G*_]. In both cases, the predictor matrix **X** is the result of concatenating the regulators and their interactions with the studied condition. In PLS1, only the potential regulators of gene g are included in **X**. In PLS2, all the potential regulators of all the G genes will constitute **X** matrix. PLS models, opposite to MLRs, take advantage of multicollinear data and can better address the high-dimensionality of multi-omics datasets as they are dimension reduction methods.

Both MLR or PLS regression models in MORE are combined with advanced variable selection strategies to properly handle high-dimensionality and multicollinearity and improve the biological interpretability of the results. These variable selection strategies are specific to each type of regression model, as described next.

### Variable selection in MORE MLR models

Omic regulators tend to act together, thus presenting highly correlated values. This multicollinearity situation hampers the estimation of MLR coefficients and the interpretation of the model, making it necessary to apply efficient variable selection methods that circumvent the problem. As detailed below, we have included in MORE two different approaches for selecting relevant regulators in MLRs: (i) a combination of a multicollinearity filter (MF) and the ElasticNet (EN) strategy, namely MF-EN, and (ii) the Iterative Sparse Group Lasso (ISGL) method.

#### (i) Multicollinearity filter and ElasticNet

The ElasticNet (EN) regularisation approach [26] for variable selection allows for having more predictors than observations in the MLR. Under a multicollinearity scenario, it is more effective than other methods, such as stepwise regression, but still presents a far from optimal performance when correlation among predictors is high. To avoid missing relevant predictors due to multicollinearity and, at the same time, improve the biological interpretation of the models, MORE performs a multicollinearity filter (MF) before defining the initial MLR equation for each gene. This MF finds groups of co-regulators (i.e. highly correlated regulators) by first creating a correlation network and next identifying highly connected groups of regulators (the detailed procedure can be found in Supplementary Note 1). By only including appropriate representative regulators from these connected groups in the model, the multicollinearity problem and the number of predictors are reduced. The correlation measurement used depends on the nature of regulator values: Pearson’s correlation for numerical regulators, Phi coefficient for binary regulators, and point-biserial correlation between numerical and binary regulators (see Supplementary Note 1).

Once the MF has been applied, MORE fits the corresponding MLR with EN penalisation for variable selection. The EN penalisation parameters (see section Supplementary Note 2) are optimised by k-fold cross-validation techniques. EN requires the predictor variables 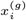 to be scaled so, given the different number of variables from each omic modality (or omic block), MORE offers three different scaling procedures [27]: auto-scaling (Equation 2), soft block-scaling (Equation 3), and hard block-scaling (Equation 4).

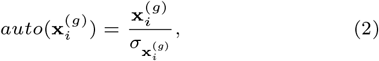

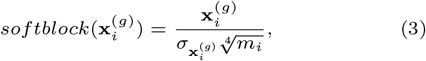

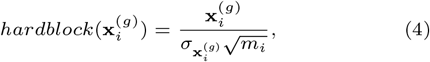

where 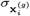 is the standard deviation of 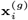 and m_*i*_ is the number of potential regulators of gene g included in the regression model belonging to the same omic block as 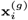.

#### (ii) Iterative Sparse Group Lasso

MORE implements another penalisation-based approach for variable selection, the Iterative Sparse Group Lasso (ISGL) penalisation [28]. As in the Sparse Group Lasso (SGL) [29] strategy, ISGL estimates a vector of regression coefficients 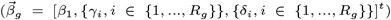 that minimises the objective function in Equation 5, which penalises the coefficients both individually and at group level.

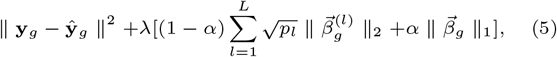

where L denotes the number of groups of predictors (regulators, in our case), p_*l*_ the number of predictors in group l, λ the shrinkage parameter, and α the parameter controlling the weight of group and individual penalisation. The ISGL method uses a reformulation of the SGL problem to optimise penalisation parameters by applying a gradient-free coordinate descent algorithm, which overcomes potential overfitting issues in high-dimensional scenarios.

The L groups of predictors in the SGL are meant to be groups of related predictors. MORE implements two different strategies to create such groups: (i) obtaining clusters of highly correlated regulators by following the MF strategy (Supplementary Note 1), or (ii) applying a Principal Component Analysis (PCA) on the predictors and assigning each predictor to the principal component (group) where it presents the highest loading [30]. As ISGL also requires scaled variables, MORE again offers the options described in Equations 2, 3 and 4.

### Variable selection in MORE PLS models

First, we introduce the notation for PLS models to explain variable selection strategies better. Equation 6 defines the matrix form of a PLS model:

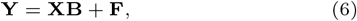

where **Y** and **X** denote the response and the predictor matrices respectively, as previously defined, **B** denotes the coefficient matrix, and **F** denotes the error matrix.

As a dimension reduction method, the idea behind PLS is to create M components or latent variables for both **Y** and **X** matrices in such a way that the covariance between **Y** and **X** components is maximised. These components are linear combinations of the original **X** variables and the projections of the n observations into these components are collected in the score matrix **T** (n × M), defined in Equation 7:

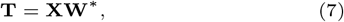

where **W**^*∗*^ = **W**(**P**^*T*^ **W**)^*−*1^ is a transformed version of the weight matrix **W**, and **P** is the loading matrix that gives the contribution of each **X** variable to each **X** component.

Similarly, the score **U** and weigth **C** matrices are defined for **Y**, such as **U** = **YC**. An iterative procedure is applied to estimate all these matrices so that the covariance between each **X** and **Y** component is maximised, and the regression coefficients matrix **B** can be computed as in Equation 8:

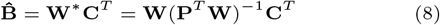

Variable scaling is also required in PLS models, so one of the three scaling approaches (Equations 2, 3, or 4) should be applied. Interestingly, MORE PLS models can handle missing data thanks to the nonlinear iterative Partial Least Squares (NIPALS) [31] algorithm applied for inferring the model.

The number of PLS components is estimated through the k-fold cross-validation procedure implemented in the ropls R package[32]. Once the PLS model is estimated, predictors are selected when having statistically significant regression coefficients and a high value of the Variable Importance in Projection (VIP) statistic. In MORE, the statistical significance of regression coefficients (H0 : b_*kl*_ = 0) can be computed either by a Jack-Knife resampling methodology or a permutation technique.

The Jack-Knife strategy removes one observation at a time; hence, n different models with n − 1 observations are fitted. Let 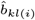 be the regression coefficient estimation for the k-th predictor and the l-th response variable when removing the i-th observation. The Jack-Knife statistic J to test H_0_ is computed as in Equation 9 and approximately follows a Student’s t distribution with one degree of freedom.

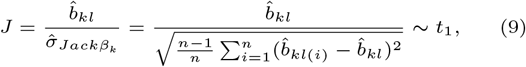

where 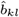 denotes the estimation of the regression coefficient for the k-th predictor (k = 1, …, p) and the l-th response variable (l = 1 for PLS1 and l = 1, …, G for PLS2).

For the permutation approach, the n observations in **Y** are randomly permuted R times. The estimated coefficients for each run 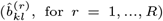 are used to create the null distribution needed to compute the statistical significance of each coefficient.

Regarding the VIP statistic (Equation 10), predictors with VIP values above a certain threshold (usually around 1) can be selected as relevant predictors in PLS models. In MORE, we combine VIP selection with either Jack-Knife or permutation strategies and select variables with significant predictors and a minimum VIP of 0.8.

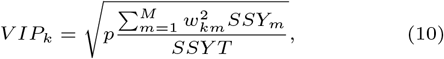

where p is the number of predictors in **X**, M is the number of PLS components, w_*km*_ is the weight of the k-th predictor in the m component, SSY_*m*_ denotes the **Y** variance explained by the m-th component, and SSY T is the total **Y** variance explained by the M components.

In summary, MORE will select predictors with a significant regression coefficient and a VIP value above the established threshold.

### MORE regulatory networks

The models and variable selection strategies implemented in MORE allow for identifying the significant regulators of each gene to build the MO-RN. As the significance of the interactions between regulators and phenotypes or conditions is also computed, we may have different regulatory situations for each phenotype or condition that, in turn, will produce a different network per condition:

- When the regulator is significant, but the interaction term is not, the regulator has the same effect on gene expression for all the conditions under study.
- When any interaction term is significant, the regulator has a different effect on gene expression for each condition.

MORE connects the significant regulators under each condition with the regulated genes in an undirected graph, where negative regression coefficients correspond to inhibitors of gene expression and positive coefficients to activators. With this information, the MO-RN for each condition is generated and can be automatically visualised through the Cytoscape platform [33]. When comparing two conditions or phenotypes (e.g. healthy and diseased), MORE can compute the differential regulatory network between both conditions to highlight the condition-specific regulations.

As the generated networks’ dimensionality may hamper the interpretability of the regulatory mechanisms, we implemented additional functionalities in MORE to assist researchers in such interpretation by connecting the relevant features in the network to biological functions or pathways. On the one hand, MORE can display the regulatory subnetwork associated with a given biological pathway. On the other hand, we implemented several options for the functional enrichment analysis of the regulatory network, as described next.

#### Functional enrichment of global regulators

In MORE, a regulator is defined as “global” when significantly regulating more than ten genes and having a network degree exceeding the third quartile of all network degrees. Global regulators are meant to play a critical role in regulatory mechanisms since their malfunction could trigger essential changes in the biological system. An important question about a given global regulator is whether it controls the regulation of a particular biological pathway; in other words, if the corresponding regulated genes are enriched in any biological pathway. To answer this question, MORE applies an Over-Representation Analysis (ORA) to the set of regulated genes versus the rest under study. In particular, we have implemented a Fisher’s exact test to perform the ORA.

However, there are significant regulators that, by nature, will never be identified as global regulators. For instance, although mutations or methylation sites rarely regulate many genes, they may still be relevant to check if genes regulated by mutations or methylation can be associated with specific functions or pathways. Therefore, MORE also offers the possibility of applying ORA to the set of genes regulated by the same omic modality.

#### Functional enrichment analysis of regulated genes

Other questions to be answered from the MO-RN network are (i) whether the highly-regulated genes (genes with many significant regulators) are involved in specific biological pathways or (ii) whether genes whose regulations change between conditions have a particular biological function. MORE addresses these two questions through a Gene Set Enrichment Analysis (GSEA) performed with the clusterProfiler R package [34]. For question (i), genes are ranked by the number of significant regulators in a given condition. For question (ii), genes are ranked according to the score S_*g*_ defined for each gene g in Equation 11) when comparing conditions A and B.

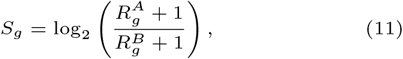

where 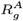 and 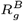 represent the number of significant regulators of gene g in conditions A and B, respectively.

MORE can also perform an ORA on the hub genes specific to a given condition.

### Benchmarking MORE

As several regression models and variable selection strategies have been implemented in MORE, we first tested them on the complete simulated data (see next section) to evaluate their performance and provide some advice on the best choices for each biological or experimental scenario.

We selected the MORE model with the best overall performance from the previous simulation study and compared it with other state-of-the-art tools for MO-RN inference. As discussed in the Introduction, we found few available tools comparable to MORE in terms of MO-RN generation with any number or types of omic modalities and for specific conditions, one of them being KiMONo [23, 24]. We also compared MORE to mixOmics [21], a reference tool in multi-omic analysis not explicitly developed for MR-RN inference.

#### KiMONo

Similarly to MORE, KiMONo is based on linear regression models to identify significant regulators of gene expression, applies SGL penalisation for variable selection where each omic modality is considered to be a group, and can (optionally) incorporate prior knowledge about potential regulations to infer the MO-RNs. KiMONo also allows the inclusion of the phenotype in the model but does not create phenotype-specific MO-RNs. KiMONo uses the igraph R package [35] to plot the inferred MO-RNs and identifies network hubs by computing node betweenness centrality. In addition, KiMONo integrates various Lasso-based strategies to handle missing data. We used version 2.0.1.0006 of KiMONo in GitHub, which depends on the hmlasso R package. This package is deprecated, so we installed it from the GitHub repository to make KiMONo work.

#### mixOmics

mixOmics incorporates different model options to analyse multi-omic data. Although none of them is directly comparable to MORE or KiMONo, we chose the sparse PLS (sPLS) [36] model for our benchmarking since it could be better adapted to the aim of this study. The sPLS methodology applies a Lasso variable selection approach to the PLS model and optimises the number of variables to be selected via cross-validation procedures. When comparing mixOmics to MORE or KiMONo, some limitations were found: (i) mixOmics does not accept prior knowledge about potential regulations. (ii) From the sPLS2 results, it is not possible to retrieve which genes are regulated by each specific significant regulator. (iii) Variables with low variability are not admitted. Considering these limitations and aiming to perform a fair comparison, we manually generated an sPLS1 model for each gene with all the omic variables as predictors to recover the specific significant regulators of each gene. We excluded regulators with a standard deviation lower than 2 to overcome the computational errors reported by the algorithm. The number of PLS components was optimised by minimising the mean absolute error (MAE). Despite the adaptations made to the mixOmics PLS model, it is still not comparable with MORE PLS1 and KiMONo because mixOmics cannot incorporate prior regulatory knowledge. Consequently, we also included MORE PLS2 in the benchmarking, which presents the same limitation. Moreover, as variable selection in mixOmics requires a grid with the potential numbers of regulators to be considered and the optimum is selected by a 2-fold cross-validation procedure with five repetitions and MAE minimisation, we adapted this grid to overcome computational limitations. For sample sizes smaller than 20 observations per group, the grid included numbers in increments of 25 regulators, 75 for 30 observations, and 150 for 50 observations.

The comparison of MORE, KiMONo and mixOmics was performed on the same simulated data but with a reduced number of regulatory omics, given the high computational requirements of mixOmics (see Data Section).

## Data

### Simulation experiments

The performance of MORE or other MO-RN methods can be defined as their capacity to detect “true” regulations. For that, we simulated a wide range of multi-omic datasets with the MOSim R package [37] where gene expression is the target “regulated” omic and the other omic modalities act as potential regulators. MOSim can simulate different omic data types and their regulatory connections to gene expression data. These simulated regulatory effects will provide the set of “true” regulators to measure the model performance. We simulated 20 different biological or experimental scenarios with MOSim. The varying factors across scenarios were:

- Sample size per phenotype (group): 5, 10, 20, 30 and 50.
- Percentage of differentially expressed genes (DEGs): 30% and 50%.
- Percentage of significant (active) regulators among the total number of potential regulators: 40% and 60%. In all cases, we established the same percentage of activator and repressor regulators but miRNA, where all significant regulators were set to be repressors.

In all of these scenarios, two phenotypes (cases and controls) and five omic modalities were simulated: gene expression (RNA-seq), transcription factor (TF) binding activity (ChIP-seq), TF expression, microRNA (miRNA) expression (miRNA-seq), DNA methylation (Methyl-seq), and chromatin accessibility (ATAC-seq). Three MO datasets were simulated for each scenario, thus resulting in the 60 MO simulated datasets we used for benchmarking.

Due to the computational cost of the mixOmics tool and its predictor variability restrictions, we used a reduced subset of the complete simulated data to compare mixOmics to MORE and KiMONo. Specifically, we only considered miRNAs and TFs as potential regulatory omics.

We first filtered out genes with an average expression of less than a count per million. Next, we applied the weighted trimmed mean of M-values (TMM) normalisation with NOISeq R package [38], voom-transformed [39] the TMM values, and computed differentially expressed genes (DEGs) between cases and controls with limma R package[40]. Genes with Benjamini and Hochberg adjusted *p* − value < 0.05 were declared DEGs.

As MOSim provides both the pairs of potential regulator-gene per omic modality and the significant regulatory pairs (let *P* be the number of significant regulations), we can consider these *P* “true” regulations as the “positive” instances and the significant regulations detected by a given method as the predicted “positives”. Therefore, we can compute a confusion matrix for each method and simulated dataset and use the F1-score metric to assess the method’s performance. In addition, as the selection of significant regulators is based on regression models for all the benchmarked approaches, the coefficient of determination R^2^ was also computed to evaluate the goodness of fit of the models obtained by each method.

### TCGA Ovarian Cancer data

A high-grade serous ovarian adenocarcinoma (HGSOC) dataset from The Cancer Genome Atlas Ovarian Cancer Cohort [41] was used to showcase MORE functionalities. The HGSOC dataset consists of 4 omic modalities measured on 292 patients: gene expression, Copy Number Variations (CNV), DNA Methylation, and microRNA expression. Data for the three first omics was downloaded from the UCSC Xena data portal [42], and normalised miRNA array data was obtained from the Broad Institute FireBrowse data portal [43].

Gene expression data was measured with RNA-seq and was already normalised and log-transformed. The NOISeq R package [38] was applied to filter out low-count genes, which resulted in keeping 20530 genes. TFs were extracted from this RNA-seq dataset and used as another regulatory layer (657 TFs). CNVs were aggregated at the gene level using the GISTIC2 method [44], resulting in CNV measurements for 24776 genes. DNA methylation was measured with Illumina microarrays, with 21666 probes, after excluding those with missing values. Methylation probes with a coefficient of variation lower than 20% were also filtered out.

We obtained the prior regulatory knowledge for MORE as follows. TF-target genes associations were obtained from the TF Link database [45]; CNVs information was already at gene level; methylation probes were associated with genes according to information provided by Illumina [46]; and target genes for the 723 miRNAs were obtained with the mirWalk web tool [47]. Specifically, we only considered associations collected in three databases: TargetScan [48], mirDB [49] and mirTarBase [50].

The phenotypes to be compared with MORE were given by the four cancer subtypes defined in [41]: *differentiated, immunoreactive, mesenchymal* and *proliferative*, also retrieved from the UCSC data portal. DEGs among subtypes were obtained with the limma R package [51] (adjusted p-value < 0.01), so 12146 DEGs were finally used as targets to generate the regulatory networks with MORE. Regulatory omic features were autoscaled, and the PLS1 model with the jack-knife resampling strategy was applied to compute the significance of PLS regression coefficients combined with a VIP value of 0.8 to select significant regulators.

## Results and Discussion

### Comparing models and strategies for variable selection in MORE

Before benchmarking MORE to other MO-RN tools, we compared the methodological options implemented in the package with different settings, as summarised in Table 1. This comparison was focused on the F1-score and considered the sample size (SS) per condition (Supplementary Figures S1 and S2). ANOVA models were used to assess significant differences in the average F1-score between models and their hyperparameters.

**Table 1.** MORE models and variable selection strategies compared on simulated data. Optimal options according to the F1-score are highlighted in bold font.

| Models | Variable Selection Methods | Parameters | Scaling type |
| --- | --- | --- | --- |
| MLR | EN + MF | r = **0.7**,<br>0.8 | **auto**<br>soft block<br>hard block |
|  | ISGL + MF |  |  |
|  | ISGL + PCA | R2 = **0.2**, 0.5,<br>0.7, 0.8 |  |
| PLS1 | Jack-Knife + VIP | $\alpha$ = 0.05<br>VIP = 0.8 | |
|  | Permutation + VIP |  |  |
|  |  | # perm = 100 |  |
| PLS2 | Jack-Knife + VIP | $\alpha$ = 0.05<br>VIP = 0.8 | auto<br>soft block<br>**hard block** |
|  | Permutation + VIP |  |  |
|  |  | # perm = 100 |  |

For single-response methods (MLR and PLS1, Supplementary Figure S1), the first expected conclusion was the superior performance of all methods for higher SS, that is, with at least 20 observations per group (ANOVA p-val < 2e − 16). Interestingly, auto-scaling worked better than block-scaling options for all methods (p-val < 2e − 16), probably because, as prior regulatory knowledge was considered, the numbers of potential regulators per omic for a given gene are not different enough in magnitude to need the block correction.

For PLS2 models (Supplementary Figure S2), as all the omic variables are included as predictors in the model, we had to reduce the omics used in this analysis only to miRNAs and TFs as MOSim generated data from over a million regulators making it unfeasible from the memory usage point of view. In this case, we got a very poor performance in terms of the F1-score, which highlights the importance of providing specific prior regulatory information for each gene to the regression models to avoid spurious associations between target features and potential regulators. Moreover, as all the omic variables are included as predictors in the PLS2 model and each omic block dimensions were still very different (130 TFs versus more than 1000 miRNAs in the simulated data used for this comparison), the hard-block scaling presented the best performance (p-val < 2e − 16), especially in low SS scenarios. Regarding variable selection, the permutation approach presented significantly better results than Jack-knife (p-val = 0.0062).

Regarding the rest of the settings for the auto-scaling option in MLR and PLS1 methods, the optimal value for the correlation threshold was r = 0.7 for both MLR+EN-MF (average F1-score 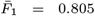) and MLR+ISGL-MF (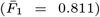), where the difference between r = 0.7 and r = 0.8 was only significant at low SS (p-val < 2e − 16, Supplementary Figure S1 A and C). In the case of MLR+ISGL-PCA (Supplementary Figure S1 B), the optimal value for the explained variance in PCA was 20% (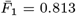). Opposite to PLS2, the Jack-Knife variable selection strategy was selected for PLS1 (Supplementary Figure S1 D,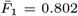), probably due to the higher dimensionality of predictor matrix in PLS2, as discussed in [52]. Finally, in Supplementary Figure S3, we introduce the F1-score results of single-response approaches across omic types using the best performing settings in each case mentioned above.

Following the optimisation of MORE strategies described above, we performed a final comparison of the resulting optimal combinations for MLR and PLS1 models in terms of F1-score, coefficient of determination (R^2^), and computational time (Figure 2) across different sample sizes per condition (SS), percentage of significant regulations among potential regulations (PSR) and percentage of DEGs (PDEG).

**Fig 2.**
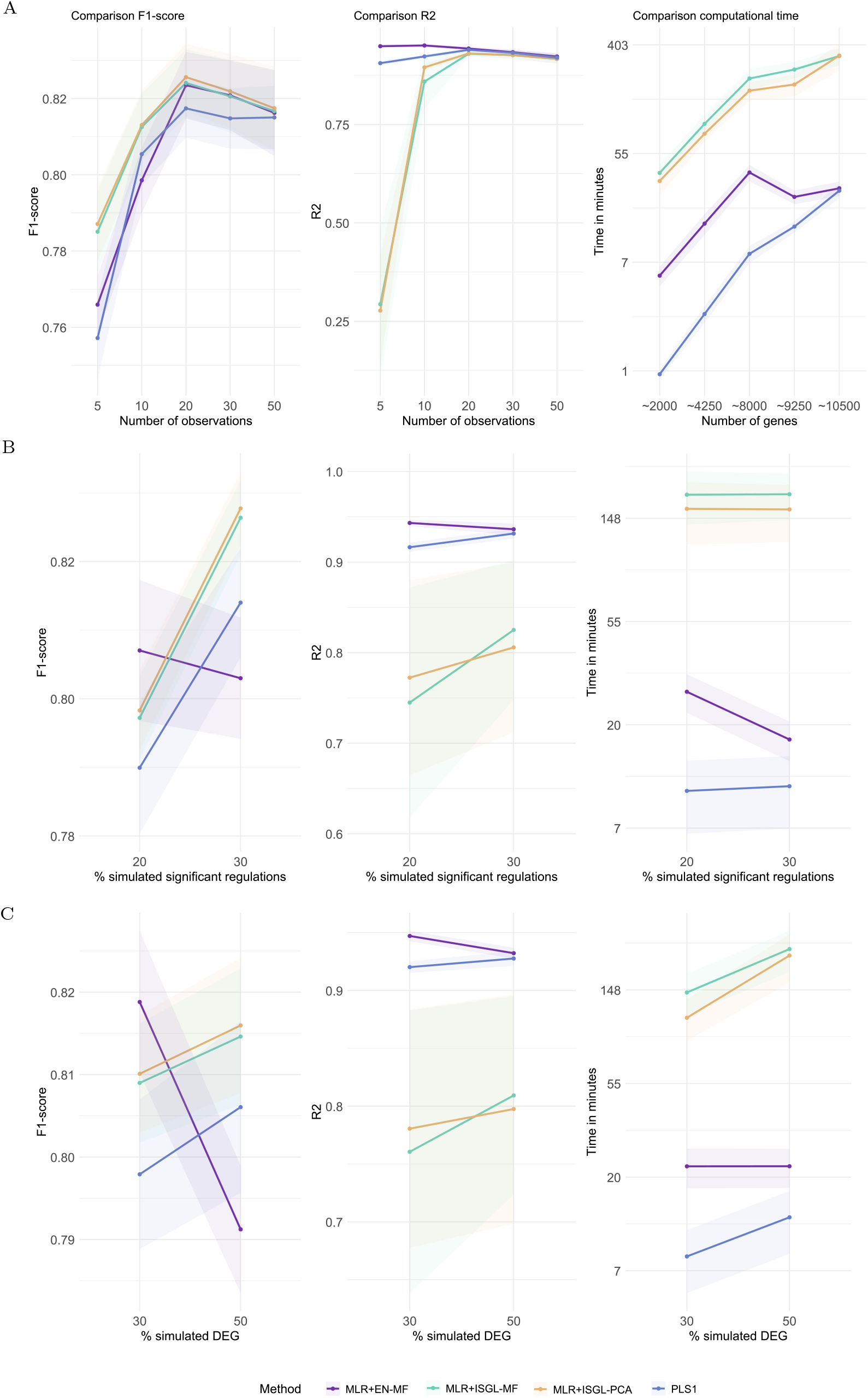
F1-score, coefficient of determination (*R*^2^), and computational time for MORE models with optimised settings. The compared models are Multiple Linear Regression with ElasticNet regularisation and multicollinearity filter (MLR+EN-MF), Multiple Linear Regression model with Iterative Sparse Group Lasso regularisation and Multicollinearity filter grouping (MLR+ISGL-MF), Multiple Linear Regression model with Iterative Sparse Group Lasso regularisation and Principal Component Analysis grouping (MLR+ISGL-PCA), and Partial Least Squares regression (PLS1) with Jack-Knife variable selection. **A)** Results across different sample sizes per condition (SS). **B)** Results for the two percentages of simulated significant regulations (PSR). **C)** Results for the two percentages of simulated differentially expressed genes (PDEG).

As expected, the SS value is critical for the performance of any methodology (p-val < 2e − 16) (Figure 2A). As previously noted, the differences in the F1-score were more pronounced (p-value < 0.0001) for low SS (n = 5 or 10) than for higher SS, where we found no significant differences among compared methods (p-val = 0.168). MLR+ISGL obtained the best performance for low SS (for both MF and PCA options), while no differences were found between MLR+EN-MF and PLS1 approaches (p-val = 0.751). The R^2^ was also affected by the SS. In this case, MLR+ISGL approaches presented lower *R*^2^ than other approaches, concluding that those approaches have high predictive performance but can not entirely explain the variability of the target omic features with the selected regulators. Regarding computational time, PLS1 was the fastest alternative, closely followed by MLR+EN-MF. The results for these two models were obtained by parallelisation with 16 cores, while the MLR+ISG strategy could not be parallelised, which explains the differences in running time.

Figure 2B shows the effect of PSR on the results. Interestingly, all methods except MLR+EN-MF present a higher F1-score and R^2^ when the PSR increases. A similar behaviour is observed when increasing PDEG (Figure 2C). The lower the PSR or PDEG, the lower the number of simulated significant regulatory relationships. Therefore, the poorer performance of MLR+EN-MF at higher levels of PSR and PDEG could indicate the difficulty of this variable selection approach when dealing with highly multicollinearity scenarios.

The benchmarking results indicate that no single method within MORE consistently outperforms the others across all scenarios. For the comparison with KiMONo, we selected the PLS1 strategy due to its minimal computational time and robust overall performance, particularly regarding R^2^. The PLS2 approach was benchmarked against mixOmics, as neither method incorporates prior regulatory knowledge. It is important to emphasise that choosing the most suitable methodology within MORE should be guided by the specific characteristics of the analysed dataset. To assist users in making this decision, we provide practical recommendations in Table 2, outlining the strengths and applicability of each approach.

**Table 2.** General recommendations to choose the most appropriate MORE settings for each experimental scenario.

|  | MLR + EN | MLR + ISGL | PLS1 | PLS2 |
| --- | --- | --- | --- | --- |
| Target omic NOT normally distributed | ✗ | ✗ | ✓ | ✓ |
| Missing data | ✗ | ✗ | ✓ | ✓ |
| High number of target features | ✓ | More time consuming | ✓ | ✓ |
| High number of regulators | ✓ | ✓ | ✓ | High memory usage* |
| Interpretation of regulations | Against a reference group |  | Against the global average |  |
| Allows for prior regulatory knowledge | ✓ | ✓ | ✓ | ✗ |
| Identifies groups of co-regulators | ✓ | ✗ | ✓ (graphically) | ✓ (graphically) |
| High multicollinearity | ✗ | ✓ | ✓ | ✓ |
| Scaling with prior regulatory knowledge for all omics | auto |  |  | block |
| Scaling without prior regulatory knowledge for any omic | block |  |  |  |

\* ~25000 regulators require more than 30GB of memory to create the model. The memory usage grows exponentially according to the number of regulators and target features included in the model. The number of phenotypes or conditions also affects.

### Benchmarking MORE to other MO-RN methods

We compared the MORE-PLS1 model (with Jack-Knife coefficient significance and auto-scaling) to KiMONo, including prior regulatory knowledge in both cases. We also compared the MORE-PLS2 model to the PLS option in the mixOmics package, as none consider prior regulatory associations. We evaluated the ability of the methods to identify the significant regulations with the F1-score, the goodness of fit of the final models with the R^2^ coefficient, and the computational efficiency (Figure 3). This benchmarking analysis was performed on the MOSim simulated datasets with only TFs and miRNAs as regulatory omics due to the computational limitations of methods that cannot incorporate prior regulatory knowledge. The results for MORE-PLS1 and KiMONo with all the regulatory omics are shown in Supplementary Material 4.

**Fig 3.**
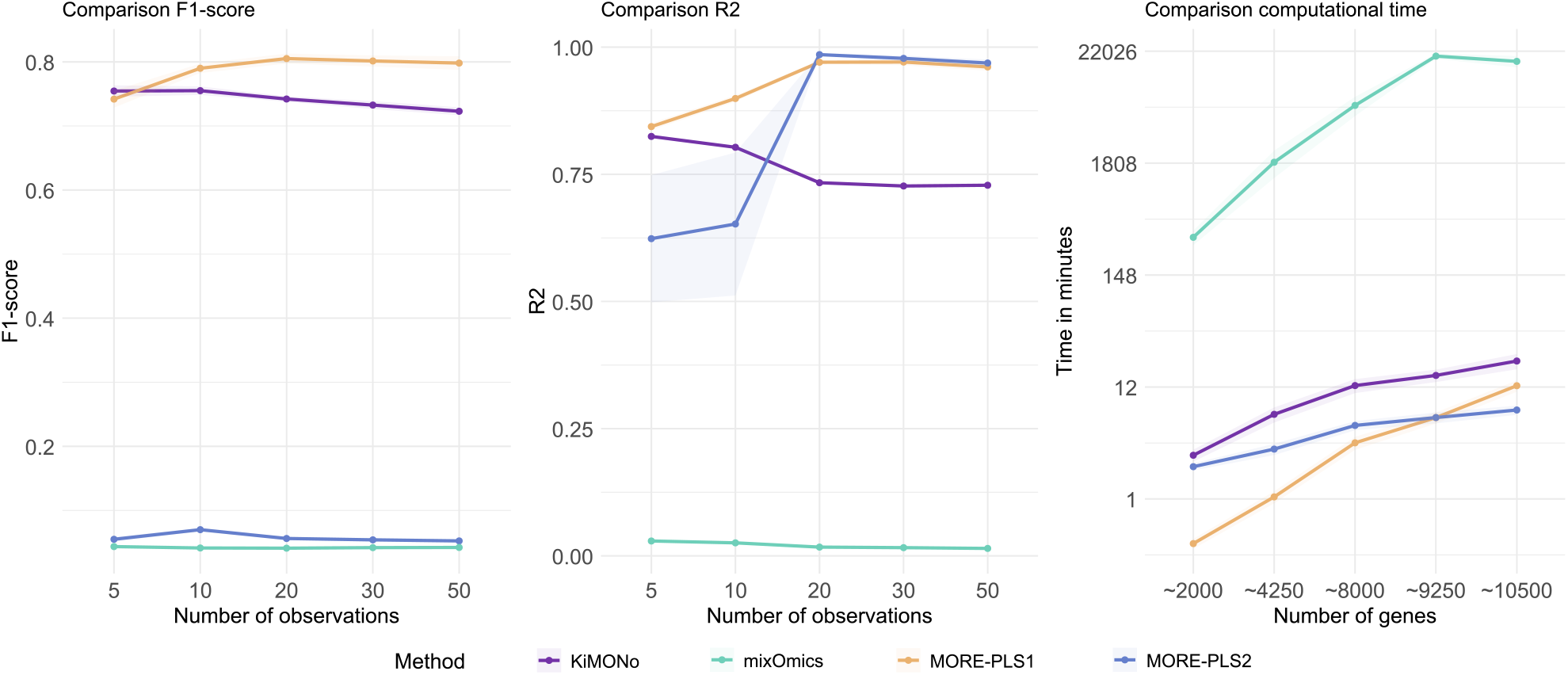
Comparison of mixOmics, KiMONo, MORE PLS1 with auto-scaling and Jack-Knife and PLS2 with hard block-scaling and permutation approaches on simulated datasets. Performance results include F1-score, coefficient of determination (R2), and computational efficiency.

The F1-score results (Figure 3) show a significant difference between KiMONo and MORE-PLS1 compared to mixOmics and MORE-PLS2. This difference is basically due to incorporating prior regulatory knowledge, highlighting that such information is essential to avoid spurious regulations, as previously noted. Interestingly, MORE-PLS1 outperforms KiMONO at all sample sizes (p-val < 2.74e − 6) but n = 5, where the difference between them is not statistically significant (p-val = 0.311). Both methods presented an overall excellent performance: MORE-PLS1 got a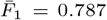, while KiMONo obtained a 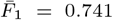. In contrast, although MORE-PLS2 significantly outperformed mixOmics (p-val = 3.2e-06), the F1-score was very low for both (below 0.06).

The superiority of MORE-PLS1 in terms of *R*^2^ (values above 0.8) is again observed for all sample sizes but *n* = 5, where the difference is not significant. Surprisingly, the *R*^2^ values for MORE-PLS2 are very high (close to 1 for higher sample sizes), while the *R*^2^ for mixOmics is below 0.1.

Finally, the most time-consuming method is mixOmics (average time of more than 9200 minutes), while MORE-PLS1, followed by MORE-PLS2, requires less time to compute the models (average time lower than 5 minutes).

When considering all the regulatory omics (Supplementary Figure S4), the differences between MORE-PLS1 and KiMONO are more evident, and MORE-PLS1 outperforms KiMONO in terms of F1-score (above 0.75 in all cases), R^2^ (higher than 0.9) and computational time. Therefore, we can conclude that MORE-PLS1 outperformed other state-of-the-art tools in all the evaluated aspects. Moreover, in the next section, we showcase the unique functionalities of MORE to aid users in interpreting the inferred MO-RNs.

### Ovarian cancer case study

We applied MORE-PLS1 to the High-Grade Serous Ovarian Cancer (HGSOC) dataset described in the Data Section, as this model presented the best performance for sample sizes of 50 observations per group, which is approximately the sample size for the four cancer subtypes we will compare in this study.

MORE fitted 11286 models, one for each gene having potential regulators in our prior regulatory information. These genes presented an average of 70.18 significant regulators. However, from now on, we will focus only on models with *R*^2^ > 0.5, corresponding to genes whose regulation was well-described. This filter reduced the number of genes in the regulatory network to 5317, with an average of 102.36 significant regulators. The percentage of genes with significant regulations by any omic modality was very similar across tumour subtypes (Figure 4A), close to 50%, and genes are regulated mainly by TFs (around 40%), closely followed by CNVs. Interestingly, the percentage of genes significantly regulated by CNVs or TFs is very similar across tumour subtypes. Still, the *proliferative* subtype presents a higher rate of regulated genes by methylation or miRNAs. Considering the initial regulatory associations provided to MORE (Figure 4B), CNVs and Methyl were naturally the omics with the lowest numbers of initial associations (10860 and 10594, respectively), but CNV presents the highest percentage of significant regulations (> 40%), followed by TFs (> 35%), and methylation (around 25%). In all omics but CNV, the percentage of significant regulations tends to increase for tumour subtypes with worse prognosis, especially for the *proliferative* subtype in methylation and TFs.

**Fig 4.**
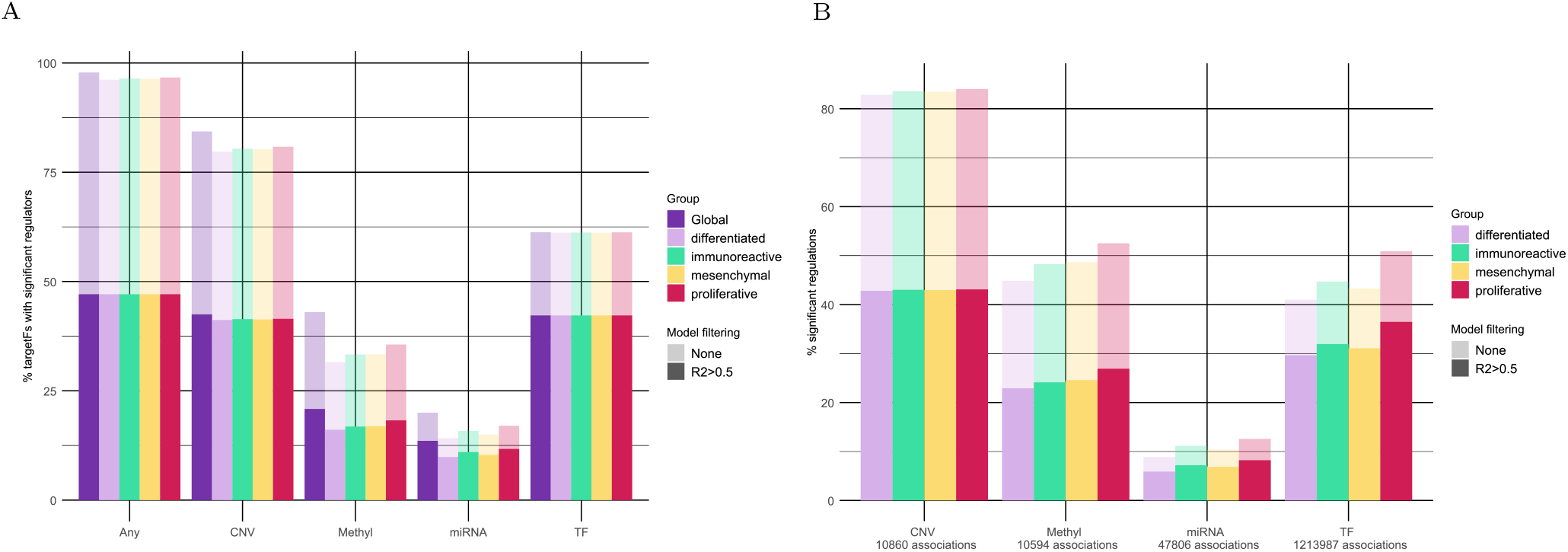
Summary of significant regulations identified by MORE-PLS1 models on the ovarian cancer data. Transparent colours correspond to results derived from all fitted models. Solid colours correspond to models with *R*^2^ *>* 0.5. A) The first set of bars shows the percentage of genes with significant regulations by any omic, globally or per tumour subtype. The following sets of bars refer to each omic modality and show the percentage of genes with significant regulations for that omic, globally or per tumour subtype. B) The percentage of significant regulations per omic over the initial potential regulations (associations) is displayed for each tumour subtype.

Figure 5A shows the complete MO-RN for the *differentiated* subtype. The interpretation of such networks becomes unfeasible due to the vast number of analysed genes and significant regulators identified by MORE models. Therefore, providing users with additional functionalities to dive into the network and extract valuable biological insights is mandatory.

**Fig 5.**
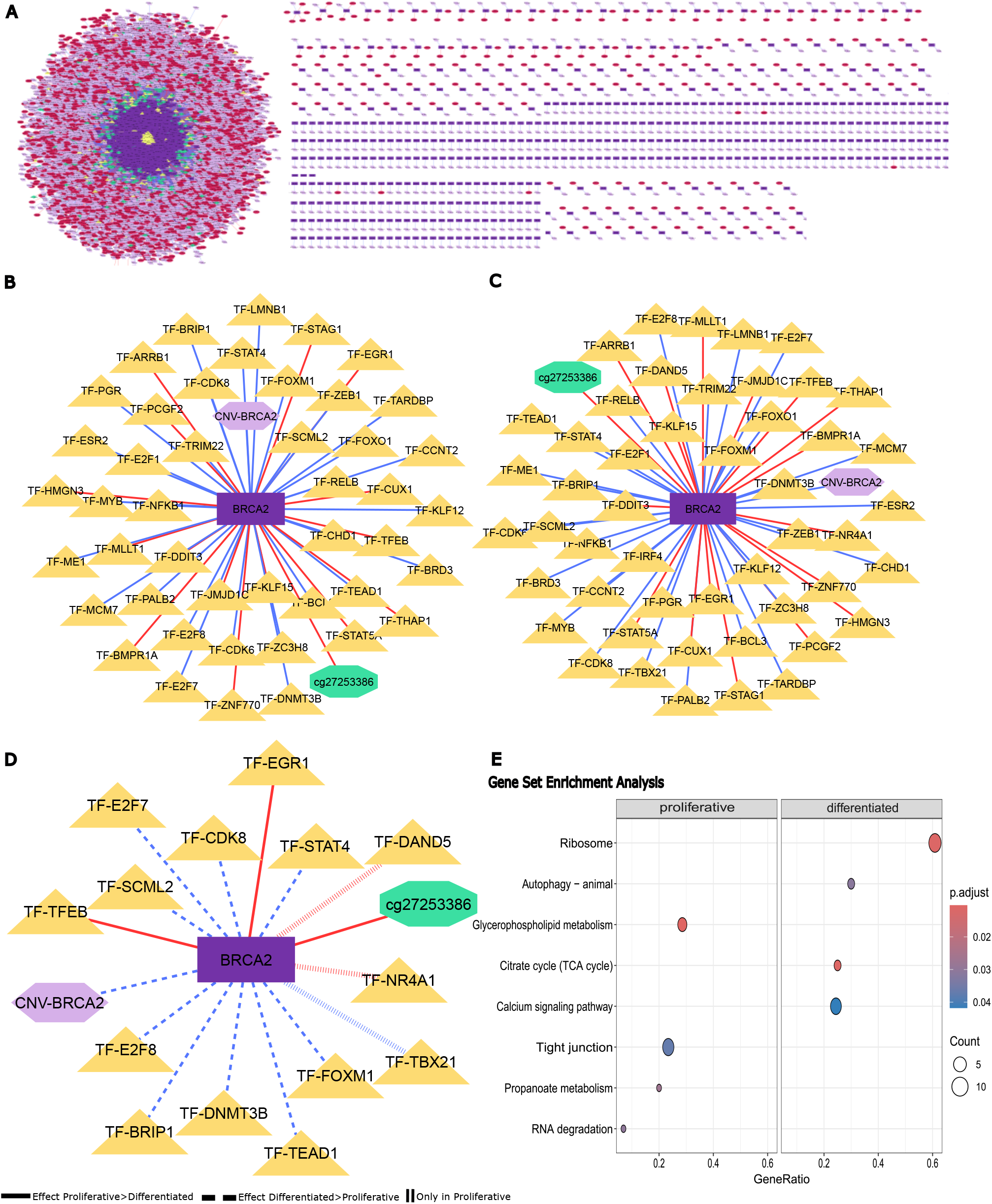
MORE results on the ovarian cancer data. **A** Regulatory network for the *differentiated* subtype. **B, C** Regulatory networks for *BRCA2* gene in the *differentiated* and *proliferative* subtypes, respectively. Red lines represent repressor effects on BRCA2 expression, and blue lines activation. **D** Differential network between *differentiated* and *proliferative* subtypes. Straight lines represent a stronger regulatory effect in the *proliferative* subtype compared to the *differentiated*. Dashed lines represent a stronger regulatory effect in the *differentiated* subtype compared to the *proliferative*. Vertical lines represent a regulatory effect only significant for the *proliferative* subtype. **E** Gene Set Enrichment Analysis results on genes ranked by the ratio between the number of MORE significant regulations in the *proliferative* subtype over the *differentiated* subtype.

A relevant question in regulatory networks is the identification of *hub genes*, defined as genes with a high number of significant regulators in the studied phenotype. Supplementary Figure S5 compares the hub genes identified by MORE for each subtype. One of the hub genes shared by all subtypes is *ZMYND8*, which has been related to a DNA damage response factor involved in regulating transcriptional responses, DNA repair activities at DNA double-strand breaks, and plays essential roles in regulating transcription during normal cellular growth [53]. Other interesting hub genes are those associated with a specific subtype since they characterise the regulation of that subtype. For example, *BRCA1* is a hub gene only in the *differentiated* s ubtype. *B RCA1* i s k nown to play a crucial role in the DNA damage repair pathway and is closely involved in the development of ovarian cancer [54, 55, 56, 57]. *BRIP1* is another interesting hub gene only present in the *differentiated* s ubtype, a lso i nvolved i n D NA damage repair, particularly in ovarian cancer predisposition [58, 59, 60]. *BRIP1* binds to *BRCA1* for homologous DNA double-strand repair and high *BRIP1* expression is related to a higher risk of recurrence and shorter disease-free survival in ovarian cancer patients treated with platinum [61], and with an increased cell proliferation [62], potentially contributing to earlier tumour recurrence [63]. The *differentiated* subtype shares the hub genes *CCNE1* and *NF1* with the *immunoreactive* subtype. Cyclin E1 (*CCNE1*) gene amplification has been reported in 15%-20% of high-grade serous ovarian tumours and has been linked to aggressive disease courses and poor outcomes [64], while the biallelic inactivation of the neurofibromatosis type 1 (*NF1*) gene has been reported as an early event in HGSOC tumorogenesis [65]. *NF1* and *MYC* are also shared with the *mesenchymal* subtype. Amplifications of MYC oncogene have been reported to occur in nearly half of HGSOC tumours and are linked to maintaining ovarian cancer cells’ oncogenic growth [66]. In the *mesenchymal* subtype, we found Fibrillin-1 (*FBN1*), which has been reported as a key factor in chemo-resistance of ovarian cancer cells by participating in the process of glycolysis and angiogenesis [67]. Finally, in the*proliferative* subtype, we found *CKB* gene. Creatine kinase B (*CKB*) is a cytosolic isoform of creatine kinase which has been reported as showing upregulated expression in several cancers at the protein level [68, 69]. This isoform has also been identified as a marker for early stages of an ovarian cancer diagnosis as its enzyme activity is significantly elevated in ovarian cancer patients, including early-stage patients [70].

For a better understanding of the differences between the *differentiated* and the *proliferative* subtypes, the two tumour subtypes with the best and worst overall survival outcome [71, 72], respectively, we performed a Gene Set Enrichment Analysis (GSEA) with MORE, where the gene score was computed as the ratio between the number of MORE significant regulations in each subtype. Functional annotation was retrieved from the Gene Ontology (GO) database [73, 74] with the AnnotationDbi Bioconductor package [75]. The functional categories more enriched in the *differentiated* subtype were “Ribosome”, “Autophagy-animal”, “Citrate cycle (TCA cycle)” and “Calcium signaling pathway”, while “Glycerophospholipid metabolism”, “Propanoate metabolism”, “Tight junction”, and “RNA degradation” were found for the *proliferative* subtype (Figure 5E). These biological functions are related to maintaining cellular structure, metabolism, energy production, and adaptation to stress, being critical in cell survival, growth, and functional regulation in multicellular organisms. Most of the altered pathways have been reported to play a crucial role in ovarian cancer; the autophagy pathway has even emerged as a therapy target in ovarian carcinoma, even if the clinical application is limited [76]. Interestingly, the Citrate cycle (TCA cycle) has been extensively reported to be an altered pathway in ovarian cancer, where the citric acid metabolite is significantly downregulated and considered to be a potential biomarker in this type of cancer[77, 78].

Other key features in regulatory networks are the *global regulators*, defined in MORE as significant regulators with many target genes, which play an essential role in the system regulation. As expected, TFs were the most abundant global regulators in all subtypes, with more than 80% representation. When focusing on the subtype-specific global regulators (Supplementary Figure S5), we found that the four subtypes shared 563 global regulators, while 15 were specific of *proliferative*, 2 of *mesenchymal*, 4 of *immunoreactive* and no specific regulators were found for the *differentiated* subtype. Remarkably, among these subtype-specific regulations, miRNAs are more prevalent: 2 of 2 in *mesenchymal*, 3 of 4 in *immunoreactive*, and 11 of 15 in *proliferative*. In the *proliferative* subtype, noteworthy miRNAs include hsa-miR-302c-3p and hsa-miR-186-5p. Interestingly, hsa-miR-302c-3p has been identified as a suppressor in the proliferation of cervical carcinoma cells, and it has been shown to inhibit the epithelial-mesenchymal transition and promote apoptosis in endometrial carcinoma cells [79]. As for hsa-miR-186-5p, this miRNA promotes the proliferation, invasion and stemness of ovarian cancer cells and plays an essential role in the regulatory effect of TUG1 on ZEB1 expression. ZEB1 has been reported to be overexpressed in multiple types of cancers and is a key transcriptional factor closely associated with epithelial-mesenchymal transition [80].

It is worth mentioning the case of the let-7 miRNA family, which has global regulators in all four subtypes. A low expression of this family has been associated with epithelial state and stemness, and these miRNAs are considered valuable predictors of HGSOC proliferation [81], as they have been directly related to migratory ability and invasiveness. We also found three members of the miR-200 family being global regulators in all the subtypes: hsa-miR-200a-3p, hsa-miR-200b-3p, and hsa-200c-3p. The miR-200 family is well-known for its implications in the diagnosis and prognosis of OC [82] and has been recognised as a master suppressor of epithelial-mesenchymal transition (EMT) [83].

Finally, it may also be interesting to study genes whose regulation was well-explained by MORE models. In particular, we focused on genes with models presenting an explained variance (R^2^) over 80%. These genes included *BRCA1, BRCA2* or *IRF2BP2*, which have been reported to play important roles in ovarian cancer [84, 85]. Figures 5B and C and D show the subnetworks of *BRCA2* for the *differentiated* and *proliferative* subtypes, respectively, while Figure 5D represents the differential subnetwork “proliferative-differentiated”. Among the differential regulators, we found BRIP1, FOXM1 and STAT4, previously reported as having relevant roles in ovarian cancer [86, 87, 88]. The three present an activator effect on *BRCA2*, stronger in the *differentiated* subtype. Notably, NR4A1 is a repressor of BRCA2 in the *proliferative* subtype, and high expression levels of this gene have been related to ovarian tumours with worse outcomes [89]. Supplementary Figure S6 shows the profiles of these four regulators and the *BRCA2* gene expression values.

## Conclusions

Understanding regulatory mechanisms from a multi-omic perspective is essential for characterising the molecular circuits driving diseases and other phenotypes of interest. However, no publicly available tools currently exist to generate phenotype-specific, multi-layered regulatory networks in an integrative manner.

In this work, we present MORE, a versatile R GitHub package designed to infer multi-omic regulatory networks. MORE enables the generation and comparison of condition-specific regulatory networks across any number or type of omics layers and facilitates the incorporation of prior regulatory knowledge into the network construction process. By leveraging regression models and advanced variable selection strategies, MORE identifies significant regulators of target features (e.g., gene expression) and provides a comprehensive suite of functionalities to aid in the interpretation of the resulting regulatory networks.

We evaluated the performance of MORE using simulated datasets, demonstrating its high accuracy and robustness, even with limited sample sizes. Additionally, we benchmarked MORE against established tools, such as KiMONo and mixOmics, in terms of identifying significant regulators, model goodness-of-fit, and computational efficiency. MORE consistently outperformed these tools across all evaluated criteria, highlighting its effectiveness and reliability.

To demonstrate its practical application, we applied MORE to an ovarian cancer dataset to study the regulation of gene expression by CNVs, transcription factors, miRNAs, and DNA methylation. We generated and compared networks for four tumour subtypes with distinct survival outcomes. The analysis revealed subtype-specific regulatory mechanisms which can explain their underlying prognostic variability. These findings emphasise the importance of capturing regulatory mechanisms not only at a global level but also in a phenotype-specific manner to better understand disease heterogeneity. This successful application of MORE highlights its utility as a powerful tool for dissecting complex regulatory networks and advancing the understanding of phenotype-specific molecular mechanisms.

## Key points

- MORE is a flexible R package designed to infer and compare condition-specific regulatory networks across multiple omics layers, addressing the lack of publicly available tools for this purpose.
- Our tool relies on a regression framework and robust variable selection techniques to identify significant regulations, ensuring accurate network inference, and can incorporate prior regulatory information.
- Evaluations on simulated data and comparisons with state-of-the-art tools demonstrate MORE’s superior performance in identifying regulators, model fit, and computational efficiency.
- MORE functionalities for downstream network analysis make it a versatile tool for interpreting complex multi-omic regulatory mechanisms.

## Authors’ contributions

S.T. and A.C. envisioned the study and conceptualised the MORE idea. S.T. supervised the work. M.C.-C. and S.T. designed and implemented a preliminary version of MORE. M.A.-M. developed and implemented the current version of MORE and performed all the analyses on simulated and experimental data. M.A.-M. and S.T. drafted the manuscript. All authors read and approved the final manuscript.

## Acknowledgment

The authors thank Carolina Monzó for her support in simulating data with the MOSim R package.

Part of the analyses were performed on the high-performance computing cluster Garnatxa at the Institute for Integrative Systems Biology (I2SysBio), I2SysBio is a mixed research centre formed by the University of Valencia (UV) and the Spanish National Research Council (CSIC).

## Data availability and software

The TCGA data is publicly available via TCGA data portal (downloaded November, 2023).

The implementation of MORE was done using R code language and it is freely available at https://github.com/Biostat-Omics/MORE/.

The results were run in a cluster with the following specifications: x86 64 architecture processor, Intel(R) Xeon(R) Gold 6230 CPU @ 2.10GHz 40c and 1500TB of RAM.

