## Supplementary information for "MORE interpretable multi-omic regulatory networks to characterize phenotypes"

#### Supplementary Note 1: Algorithm for multicollinearity filter

**Algorithm 1** Multicollinearity filter structure

```
procedure
    Compute the correlation for each pair of regulators:
    Continuous regulators → Pearson's coefficient
    Binary regulators → Phi's coefficient
    Binary vs Continuous regulator → Point biserial coefficient
    Generate networks connecting regulators with a correlation higher than the threshold specified by the user
    Obtain connected components of the network
    for each connected component do
        if only one regulator then
            Include in the MLR model
        else
            while there are connected components do
                if it is a fully connected graph then
                    Take as representative the one with highest correlations
                else
                    Take as representative the one with highest degree (in case of a tie, the one with highest correlations)
                    Remove from the network the associated regulators
                end if
            end while
        end if
    end for
end procedure
```

To compute correlation between regulators $i$ and $j$, with vectors of values $\mathbf{x}_i^{(g)}$ and $\mathbf{x}_j^{(g)}$, respectively, we used the following correlation measurements:

**Pearson's correlation coefficient**

$$\rho_{\mathbf{x}_i^{(g)},\mathbf{x}_j^{(g)}} = \frac{cov(\mathbf{x}_i^{(g)},\mathbf{x}_j^{(g)})}{\sigma_{\mathbf{x}_i^{(g)}} \cdot \sigma_{\mathbf{x}_j^{(g)}}} \tag{1}$$

where $cov$ denotes the covariance and $\sigma_{\mathbf{x}_i^{(g)}}$ and $\sigma_{\mathbf{x}_j^{(g)}}$ the standard deviation of $\mathbf{x}_i^{(g)}$ and $\mathbf{x}_j^{(g)}$ numerical omic regulators, respectively.

**Phi's correlation coefficient**

$$\phi_{\mathbf{x}_i^{(g)},\mathbf{x}_j^{(g)}} = \frac{(ad - bc)}{\sqrt{(a+b)(c+d)(a+c)(b+d)}} \tag{2}$$

where $a$, $b$, $d$ and $c$ are the counts of the contingency table, and $\mathbf{x}_i^{(g)}$ and $\mathbf{x}_j^{(g)}$ binary omic regulators.

**Point biserial correlation coefficient**

$$r_{\mathbf{x}_i^{(g)},\mathbf{x}_j^{(g)}} = \frac{\overline{\mathbf{x}}_{i1}^{(g)} - \overline{\mathbf{x}}_{i0}^{(g)}}{\sigma_{\mathbf{x}_i^{(g)}}}\sqrt{pq}, \tag{3}$$

where $\mathbf{x}_i^{(g)}$ is a numerical regulator and $\mathbf{x}_j^{(g)}$ a binary regulator, $p$ is the proportion of 1's in the binary regulator, $q = 1 - p$ the corresponding proportion of 0's, $\overline{\mathbf{x}}_{i1}^{(g)}$ and $\overline{\mathbf{x}}_{i0}^{(g)}$ are the conditional means of the numerical regulator when the binary regulator is 1 or 0, respectively and $\sigma_{\mathbf{x}_i^{(g)}}$ is the standard deviation of the numerical regulator $\mathbf{x}_i^{(g)}$.

As correlation calculation can sometimes lead to spurious relationships, we also included the full-order partial correlation coefficient:

$$\rho_{\mathbf{x}_i^{(g)}\mathbf{x}_j^{(g)}\cdot\mathbf{X}\setminus\{\mathbf{x}_i^{(g)},\mathbf{x}_j^{(g)}\}} = -\frac{p_{\mathbf{x}_i^{(g)}\mathbf{x}_j^{(g)}}}{\sqrt{p_{\mathbf{x}_i^{(g)}\mathbf{x}_i^{(g)}}\,p_{\mathbf{x}_j^{(g)}\mathbf{x}_j^{(g)}}}}, \tag{4}$$

where $(p_{\mathbf{x}_i^{(g)}\mathbf{x}_j^{(g)}}) = \Sigma^{-1}$ is the inverse of the correlation matrix computed with the coefficients described above.

#### Supplementary Note 2: ElasticNet variable selection strategy

The ElasticNet (EN) variable selection method [1] estimates a vector of regression coefficients $\vec{\beta}_g = [\beta_1, \{\gamma_i, i \in \{1, ..., R_g\}\}, \{\delta_i, i \in \{1, ..., R_g\}\}]^t$ such as the following objective function is minimised:

$$\| \mathbf{y}_g - \hat{\mathbf{y}}_g \|^2 + \lambda[(1-\alpha) \| \boldsymbol{\beta}_g \|^2 + \alpha \| \boldsymbol{\beta}_g \|_1], \tag{5}$$

where $\lambda$ is the shrinkage parameter and $\alpha$ controls the weight of Ridge regression [2, 3] and Lasso [4] methods in the EN model. A k-fold cross-validation procedure is applied to optimise these two penalisation parameters. The number of folds $k$ is set according to the sample size (varying from 5 to 10), and a leave-one-out cross-validation is applied when the sample size is lower than 50.

### Supplementary Material 3: Comparison of results on simulated data for MORE models under its various settings

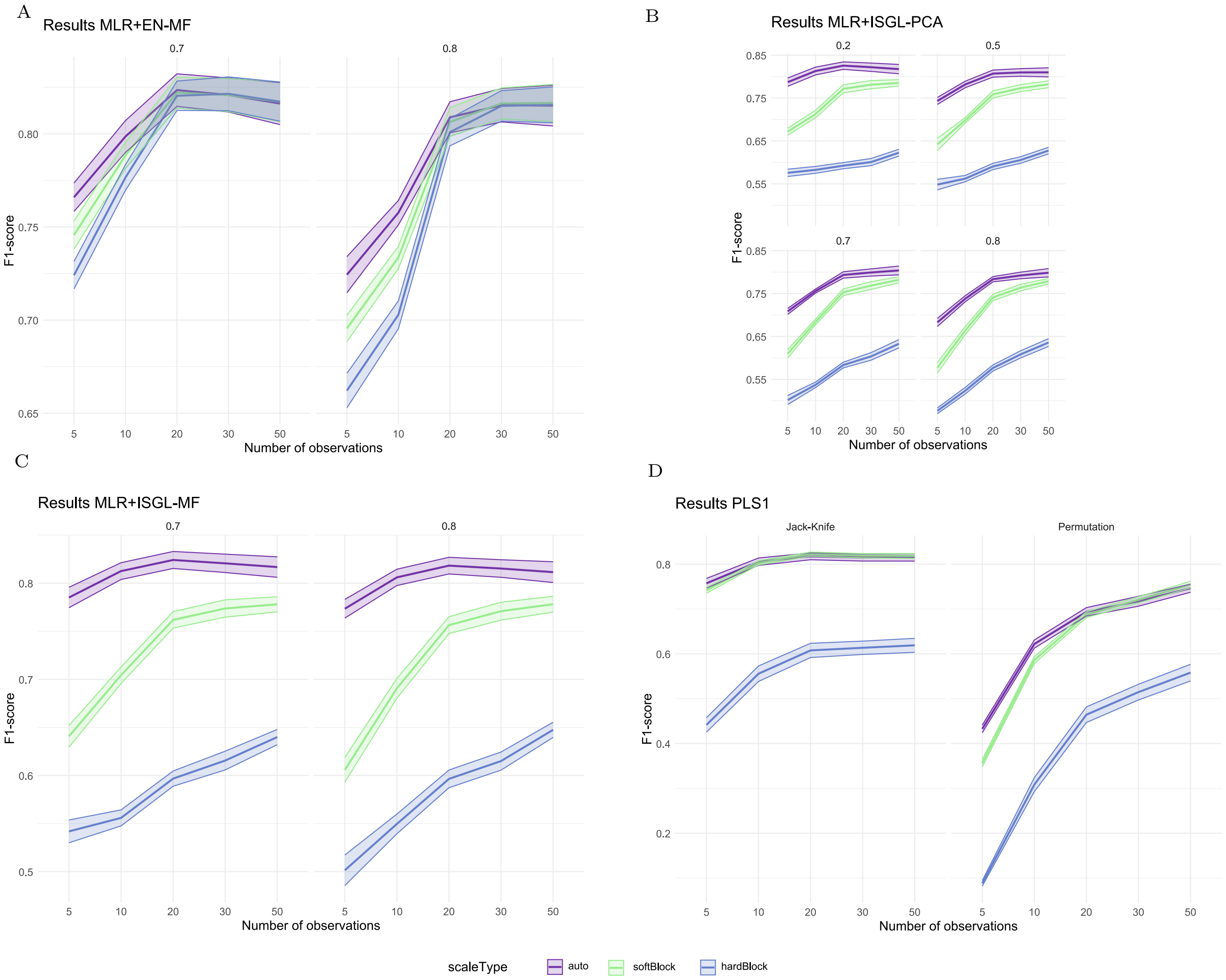


**Fig. S1.** F1-score results for different settings of MORE methods. The results are compared for different scaling approaches: auto-scaling, soft block-scaling and hard block-scaling. The straight lines show the mean, while the bandwidth represents the confidence interval of the results. **A)** F1-score results for Multiple Linear Regression models with ElasticNet regularisation and Multicollinearity filter for different correlation thresholds for the filtering: 0.7 and 0.8. **B)** F1-score results for Multiple Linear Regression models with Iterative Sparse Group Lasso regularisation and Principal Component Analysis grouping approach for different thresholds for the percentage of variability to explain: 0.2, 0.5, 0.7 and 0.8. **C)** F1-score results for Multiple Linear Regression models with Iterative Sparse Group Lasso regularisation and Multicollinearity filter grouping approach for different correlation thresholds for the filtering: 0.7 and 0.8. **D)** F1-score results for Partial Least Squares models for different approaches for estimating statistical significance: Jack-Knife and Permutation resampling.

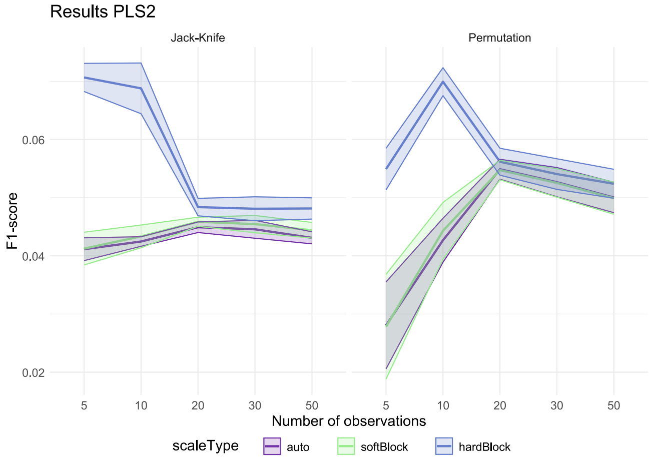


**Fig. S2.** F1-score results for Partial Least Squares 2 models for different approaches for the coefficient significance estimation: Jack-Knife and Permutation resampling. In this case, a unique model is created for all genes without prior regulatory knowledge taking only miRNAs and TFs as potential regulatory omics.

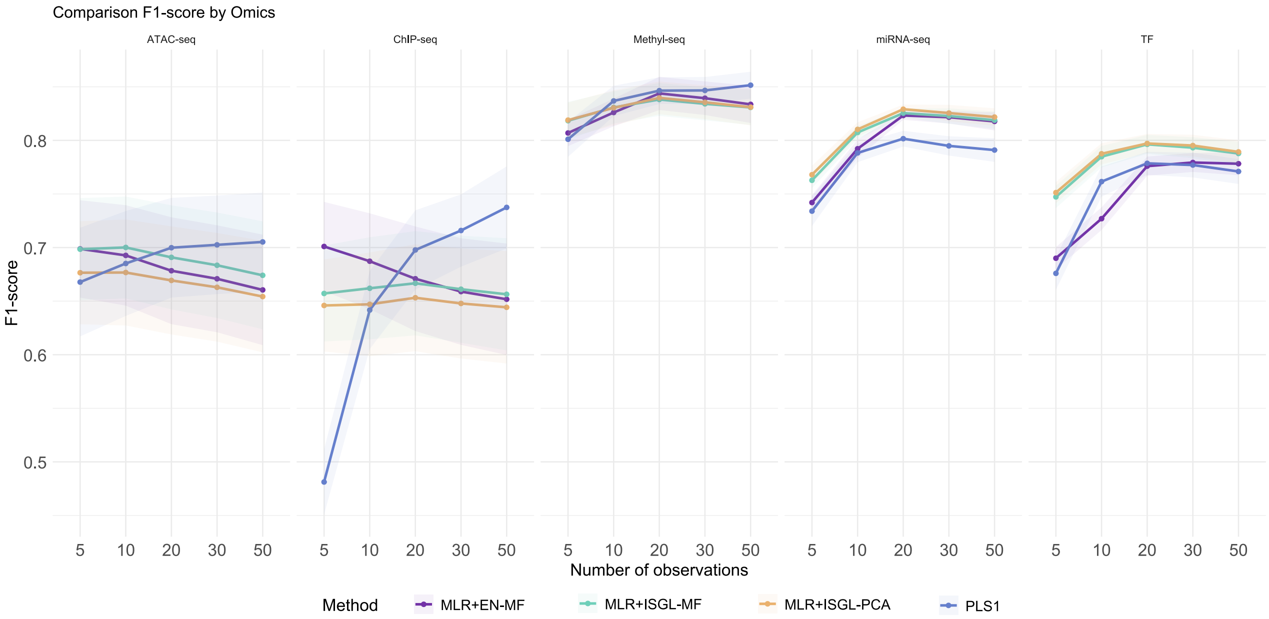


**Fig. S3.** F1-score of the best settings of the different MORE methodologies. Results per number of observations and omic analysed. Results of the following models: Multiple Linear Regression model with ElasticNet regularisation and multicollinearity filter with a threshold of 0.7 for the filtering, Multiple Linear Regression model with Iterative Sparse Group Lasso regularisation and Multicollinearity filter grouping with a correlation threshold of 0.7, Multiple Linear Regression model with Iterative Sparse Group Lasso regularisation and Principal Component Analysis grouping for 20% threshold for the percentage of variability to explain, and Partial Least Squares regression with Jack-Knife methodology for the estimation of the significance of the coefficients of the regression model. In all cases, the scaling used was auto-scaling.

We compared the best-performing settings for each methodology (from Supplementary Figure S1) based on the F1-score across omic types. The selected methodologies were: MLR with EN regularisation and MF with a threshold of 0.7 for the filtering, MLR with ISGL regularisation and MF grouping with a correlation threshold of 0.7, MLR with ISGL regularisation and PCA grouping for 20% threshold for the percentage of variability to explain, and PLS1 with Jack-Knife methodology for the estimation of the significance of the coefficients of the regression model. In all cases, the scaling used was auto-scaling.

ChIP-seq presented the worst results, followed by ATAC-seq, likely due to ChIP-seq being simulated as a binary omic with inherently different data compared to others. MLR approaches showed more stable results across sample sizes compared to PLS1, however PLS1 performed significantly at higher sample sizes. Methyl-seq presented the best F1-score for all methodologies, which was surprising since $\beta$ values were considered, so a priori, these regulators were the ones that showed the least variability. For miRNA-seq and TF, MLR+ISGL approaches performed better, but overall, they presented an expected behaviour with F1-scores increasing as sample size grew.

#### Supplementary Material 4: Comparison of results on simulated data between MORE's best-performing method and KiMONo

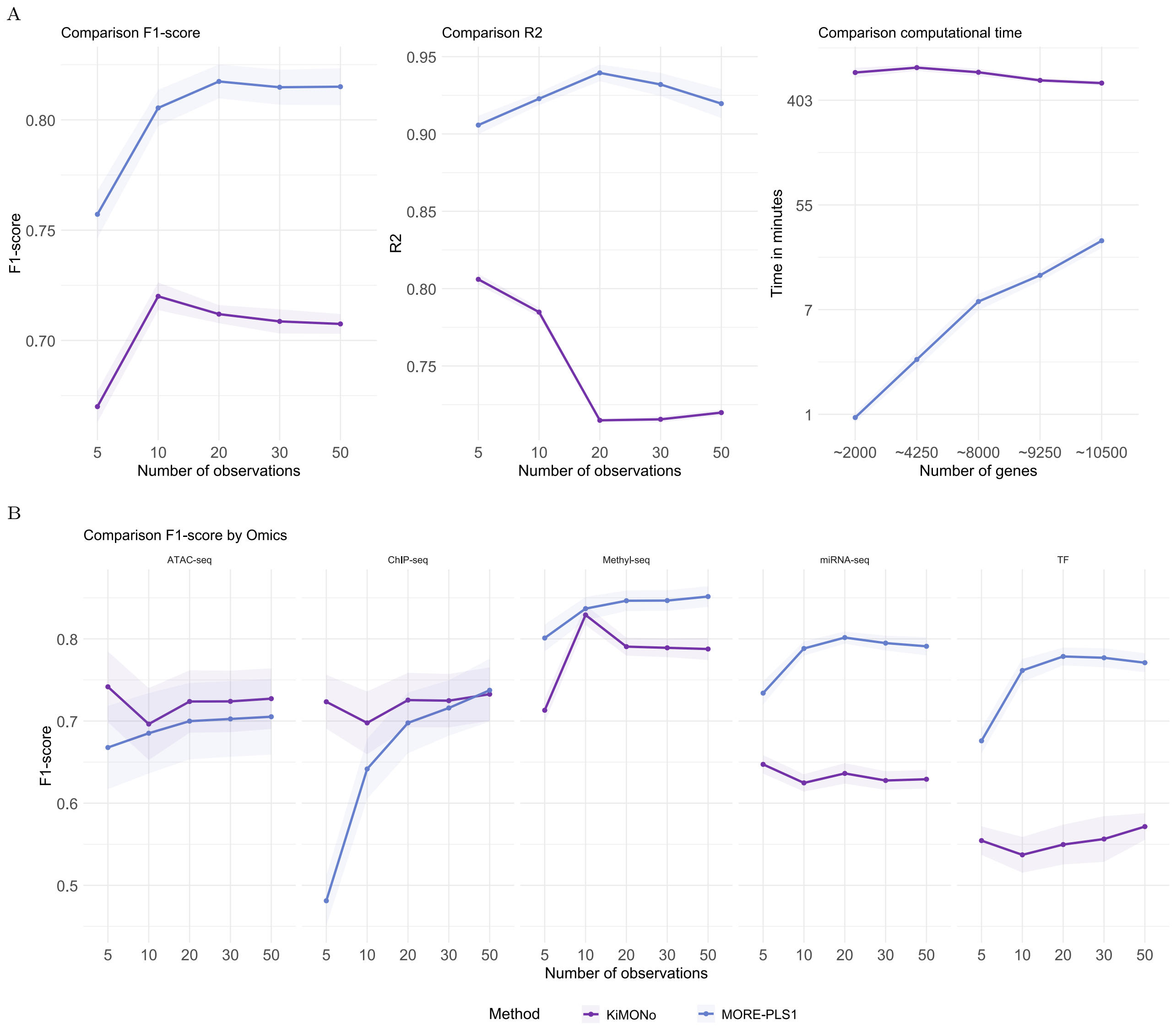


**Fig. S4.** Results of MORE's best approach (PLS1+Jack-Knife+auto-scaling) compared to KiMONo. **A)** Comparison of F1-score, $R^2$ and computational efficiency. **B)** F1-score results through the different omics.

#### Supplementary Material 5: MORE use-case in HGSOC

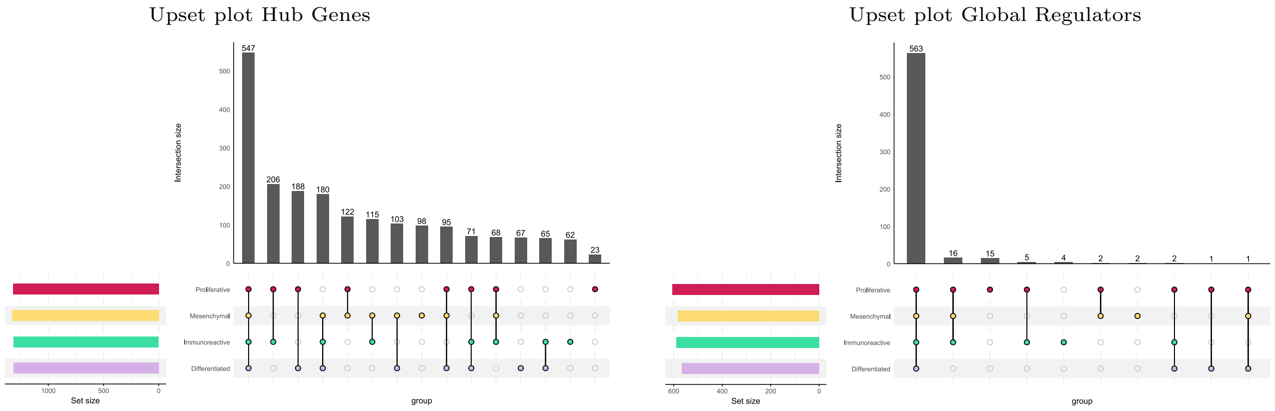


**Fig. S5.** Number of hub genes and global regulators shared by the different cancer subtypes

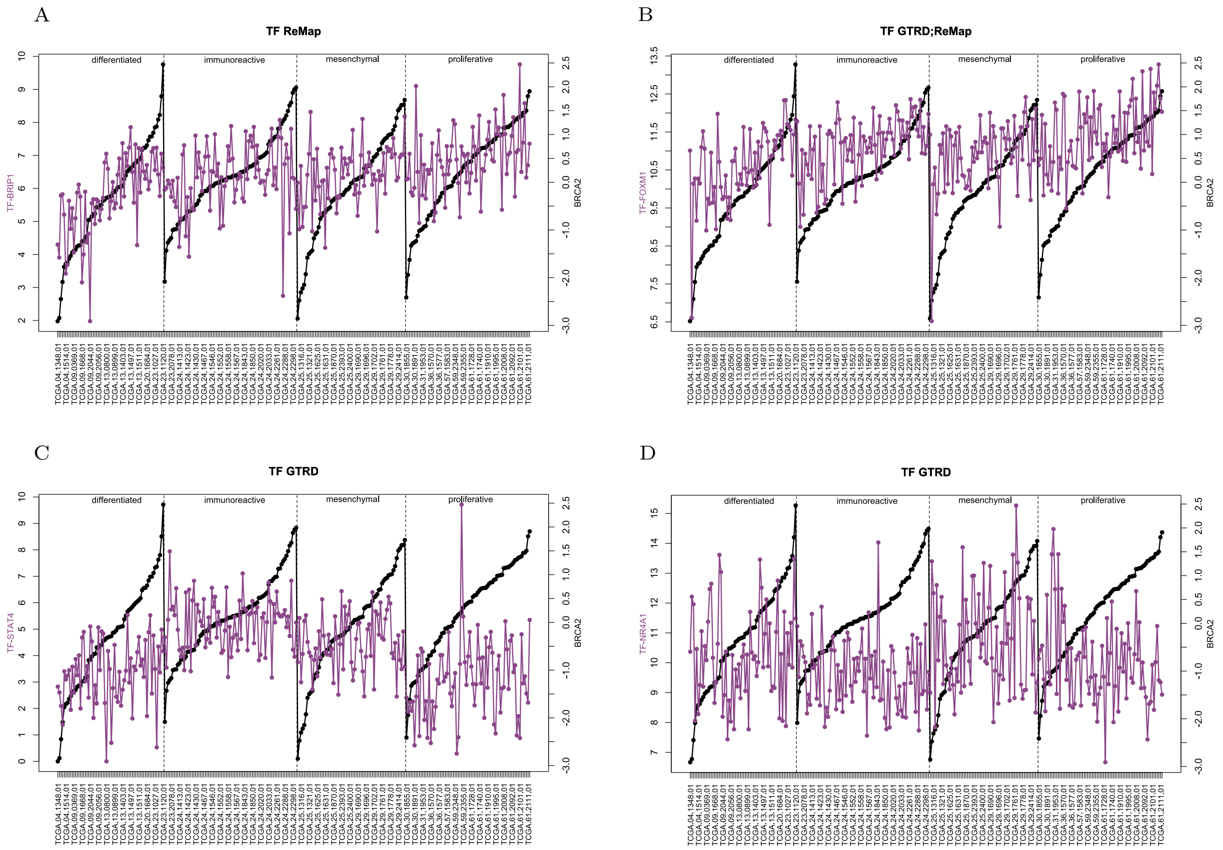


**Fig. S6.** Gene versus regulator expression profiles for the regulators that significantly regulated BRCA2 and which presented a different behaviour in *differentiated* compared to the *proliferative* subtype

Figures S6A, S6B and S6C show, as Figure 5D already presented, how TFs *BRIP1*, *FOXM1* and *STAT4* presented an activator effect on *BRCA2* gene expression. What is more, in these figures, it could be seen how the relation between the expression of the regulators and the gene expression is tighter in *differentiated* compared to *proliferative* subtype, which supports the findings of MORE. In the case of TF *NR4A1*, MORE found a significant regulation only for the *proliferative* subtype, which presented a repressor effect on the gene expression of *BRCA2*. Once again, S6D showed how the *differentiated* subtype presented almost a random profile for the regulator while the *proliferative* samples with lower gene expression presented higher expression on the regulator. These results support the results found by MORE in the first place.
